# TURNOVER OR INTRASPECIFIC TRAIT VARIATION: EXPLAINING FUNCTIONAL VARIABILITY IN A NEOTROPICAL ANURAN METACOMMUNITY

**DOI:** 10.1101/2020.03.04.977397

**Authors:** Diego Anderson Dalmolin, Alexandro Marques Tozetti, Maria João Ramos Pereira

## Abstract

Trait variation across environmental gradients results from two processes: intraspecific variation (ITV) and turnover. Tadpoles are known to exhibit phenotypic plasticity in several traits in response to the environment, resulting from intra or interspecific variation. Here we evaluate patterns of intraspecific variation in functional traits of adult anurans (head shape, eye size and position, limb length and body mass) and their relationship with environmental variables in an anuran metacommunity in southern Brazil. From anurans sampled from 33 ponds, we decomposed trait variation into ITV and turnover and modelled trait-environment relationships. We predict that the contribution of ITV and turnover to trait variation and trait-environment relationships should vary according to the preferred habitat of the species and the analysed traits. Intraspecific variation accumulated the highest rate of trait variation for arboreal species, while interspecific variation was greater for aquatic-terrestrial species and for the whole set of species. The contributions of turnover and ITV to shifts in community mean trait values were similar between traits, but differed between species sets. Depth, distance between ponds, area of *Pinus* surrounding the ponds, and types of pond vegetation and substrate strongly influenced trait variation, but their relative contribution depended on the analysed traits and species sets. The great contribution of ITV for head shape and eye size and position suggests the existence of intraspecific adaptations to microhabitats, while turnover dominance in the variation of body mass and limb length suggests differences in dispersal and trophic segregation between species.

## 1 INTRODUCTION

Functional-based approaches quickly gained prominence in community ecology. Functional traits are excellent tools for assessing the influence of ecological predictors (environmental and spatial) on community assembly patterns as well as of processes of colonization and species maintenance in different environments and geographical areas (McGill et al., 2006; Meynard et al., 2011; Tonkin et al., 2016). Ecologists now have a wide variety of functional diversity metrics - see the reviews by Villéger et al. (2010) and Mouchet et al. (2010). However, the vast majority of studies is based on assessments focusing almost entirely on mean attribute values (Luo et al., 2016). Consequently, the observed patterns of functional dissimilarity between communities were mostly attributed to interspecific trait variation, neglecting valuable information provided by intraspecific variation (McGill et al. 2006; Dahirel et al., 2017; Fajardo and Siefert, 2018).

Recent studies suggest that the quantification of intraspecific trait variation patterns (henceforth ITV) is paramount, because it allows a more accurate view of the functional organization of communities and patterns of species coexistence (Bolnick et al., 2011; Violle et al., 2012; Kraft et al., 2015). It has been demonstrated that ITV for several groups of organisms across different environmental gradients is high - e.g. trees, Jung et al. (2010), Fajardo and Siefert (2018); invertebrates, Rudolf and Rasmussen (2013), Dahirel et al. (2017); and tadpoles, Zhao et al. (2017), Jordani et al. (2019) – strongly contributing to the increase of alpha and beta diversities (e.g. Spasojevic et al., 2016). Changes in environmental conditions lead to the expression of phenotypic plasticity in several traits (e.g. morphological, behavioural and life history) within and between populations (Mouillot et al., 2013), and, for that reason, ITV is considered the main link between mechanistic processes and community structure (Lepš et al., 2011; Siefert et al., 2015).

Community-level trait responses to niche-based processes are driven by a combination of turnover and ITV (Lepš et al., 2011). In situations where ITV is the dominant process, attribute values will vary across the environmental gradient without modification in species composition. In the opposite case, i.e., when turnover dominates, it is the species composition that varies across the gradient, with little or no change in attribute values (Lepš et al., 2011; Violle et al., 2016). Most studies published so far detected an alternation in the dominance of both processes in the ITV patterns between different types of traits along environmental gradients (e.g. Volf et al., 2016; Derroire et al., 2018; Fajardo and Siefert, 2018). Changes in the balance between those processes leading to ITV may drive the occurrence of individuals in certain environments because they facilitate or prevent the coexistence between organisms of the same species (Turcotte and Levine, 2016). Consequently, the number of individuals from each species that overlap in niche space will decrease (Cloyed and Eason, 2017).

Anurans are important components of freshwater ecosystems: they represent a substantial part of the biomass of those environments and usually occupy several positions of the trophic web (Gibbons et al., 2006; Wells, 2010; Rowland et al., 2015). The structure of anuran communities in ponds tends to respond to frequent, and often unpredictable, environmental changes - such as changes in water depth, presence and abundance of aquatic vegetation and hydroperiod (Wellborn et al., 1996; Urban, 2004; Richter-Boix et al., 2007; Werner et al., 2007). In fact, pond dynamism prompted by changes in local environmental conditions leads to different degrees of trade-offs (e.g. growth and survival) and high rates of trait diversification and specialization in resident species (e.g. body size and larval development rates; Wellborn et al., 1996; Schemske et al., 2009; Schalk et al., 2017). This pattern may be quite intense in anurans, as they have a biphasic life cycle, highly permeable skin and are highly sensitive to environmental changes during most part of their life-cycle (Niemi and McDonald, 2004). Despite the scarcity of information for adults, previous studies have shown high phenotypic plasticity in tadpoles for various attributes across different ecological gradients (Michel, 2011; Marques and Nomura, 2015; Marques et al., 2019).

In this study we used a trait-based approach to evaluate patterns of intraspecific variation and their relationship with local environmental predictors in an anuran metacommunity of 33 ponds located in the southernmost coastal region of Brazil. We test the hypothesis that species ITV and turnover processes influence trait variation patterns across environmental gradients. We expected traits related to habitat (and microhabitat) exploration and foraging – mostly those related to the trophic and functional positioning of individuals (head shape, eye position and eye size), that are promoters of the stabilization of niche differences between species – to be less variable within species in comparison to those related to physiological dispersion and control (body mass and relative limb length; Turcotte and Levine, 2016, Jordani et al., 2019) which should be more influenced by species turnover.

In addition, we evaluate the existence of possible relationships between different sets of environmental predictors and the patterns of variation of functional traits. Trait-environment relationships should be more evident between similar ecological groups, which may respond to similar environmental predictors. We expect traits related to habitat exploration/foraging to be driven by variables related to structural variation of ponds (Prado et al., 2014; Leão-Pires et al., 2018; Lima et al., 2019). We also expect traits related to dispersal and physiological control (water conservation hypothesis - Amado et al., 2019) to be associated with the geographic distance between ponds, variables reflecting hydroperiod and matrix surrounding the ponds (da Silva et al., 2012, Machado et al., 2012; Prado et al., 2014; Queiroz et al., 2015; Tavares and da Silva, 2019).

## 2 MATERIAL AND METHODS

### 2.1 Ethics statement

We obtained the collecting permits from Instituto Chico Mendes de Conservação da Biodiversidade (ICMBio) (licence 55409). Our sampling did not involve any endangered or protected species. We restricted amphibian manipulation in the field to the minimum necessary (see section 2.3); specimens collected were identified to species level, measured and immediately released after these procedures in the same pond/site where they were captured.

### 2.2 Study Area

This study was carried out in Lagoa do Peixe National Park (PNLP; 31°02_-31°48_S; 50°77_-51°15_W; figure 1), one of the two Ramsar sites in southern Brazil (RAMSAR, 2018). With a length of 64 km and an average width of 6 km, the PNLP comprises over 34,000 hectares of protected wetlands, integrating the Coastal Plain of the State of Rio Grande do Sul (Maltchik et al., 2003). The climate is subtropical humid, and temperatures range between 13 °C and 24 °C with annual average of 17.5 °C. The mean annual precipitation varies between 1200 and 1500 mm (Tagliani, 1995).

**Figure 1:**
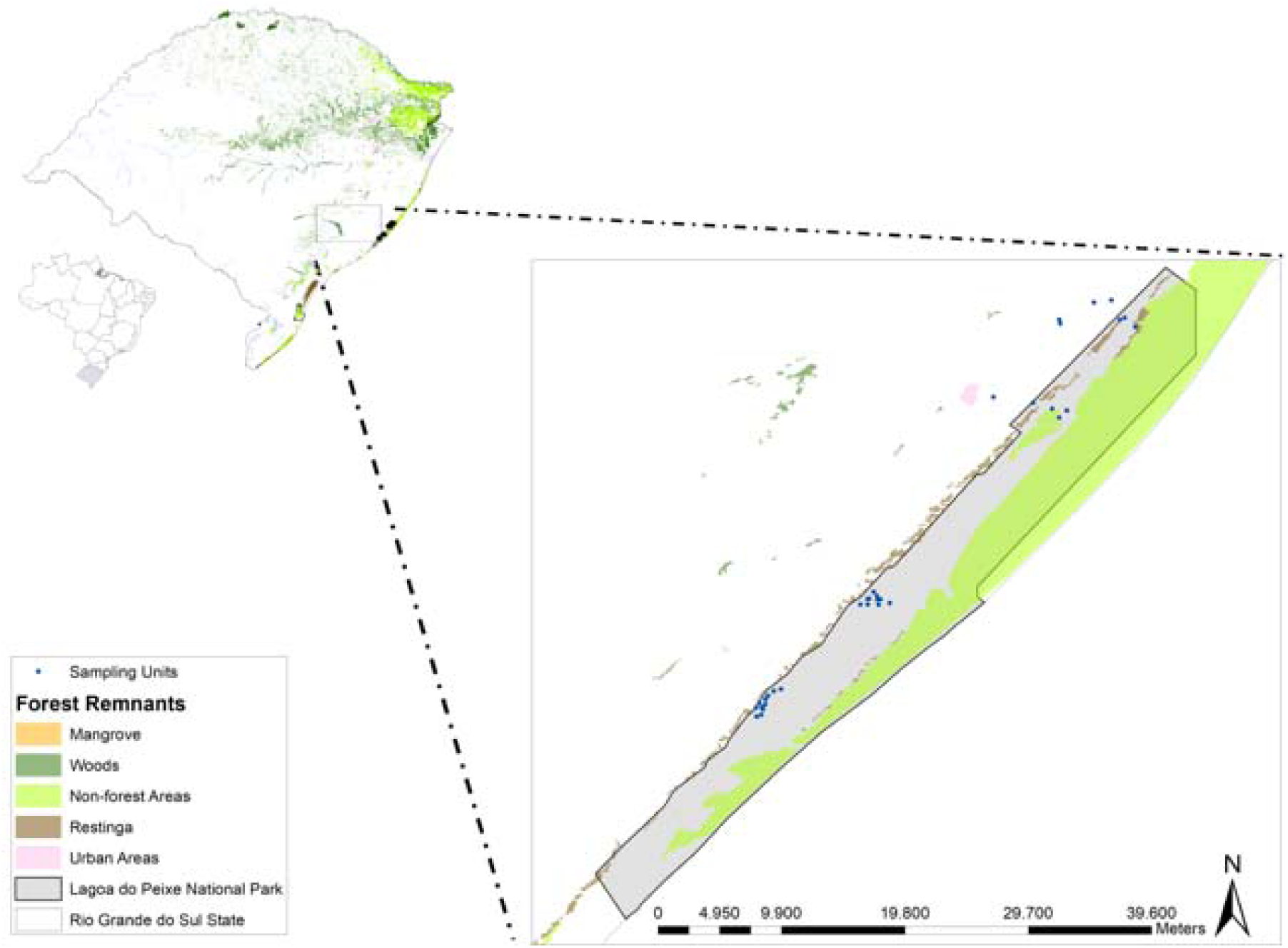
Study area at Lagoa do Peixe National Park, southern Brazil. Sampled ponds are represented by blue circles (N = 33).

### 2.3 Anuran surveys and trait measurement

We sampled adult anurans in 33 ponds throughout the study area (Fig. 1). We selected the ponds based on biotic and abiotic characteristics (see Table S1) and on the distance between ponds to guarantee spatial independence. Distance between ponds ranged from 0.7 to 39 km. Sampling was performed monthly, from October 2016 to March 2017. We used both calling surveys and active search at breeding habitats to find the individuals in each pond (Scott and Woodward, 1994). Samplings were carried out from 6 p.m. to 0 a.m; monthly effort per pond was of 1 hour, totalling 6 hours of sampling per pond.

We evaluated five functional traits in five individuals per species per pond: head shape; eye position; eye size; relative length of limbs; body mass. We chose these traits based on perceived importance for determining habitat use and species resilience, and they were characterized according to the metrics presented in Figure 2 and Table 1. To avoid reassessment of the same individual in the same sampling period, we placed all individuals captured in containers for the duration of the sampling of the respective pond and then released them at the exact same site of capture.

**Table 1.**
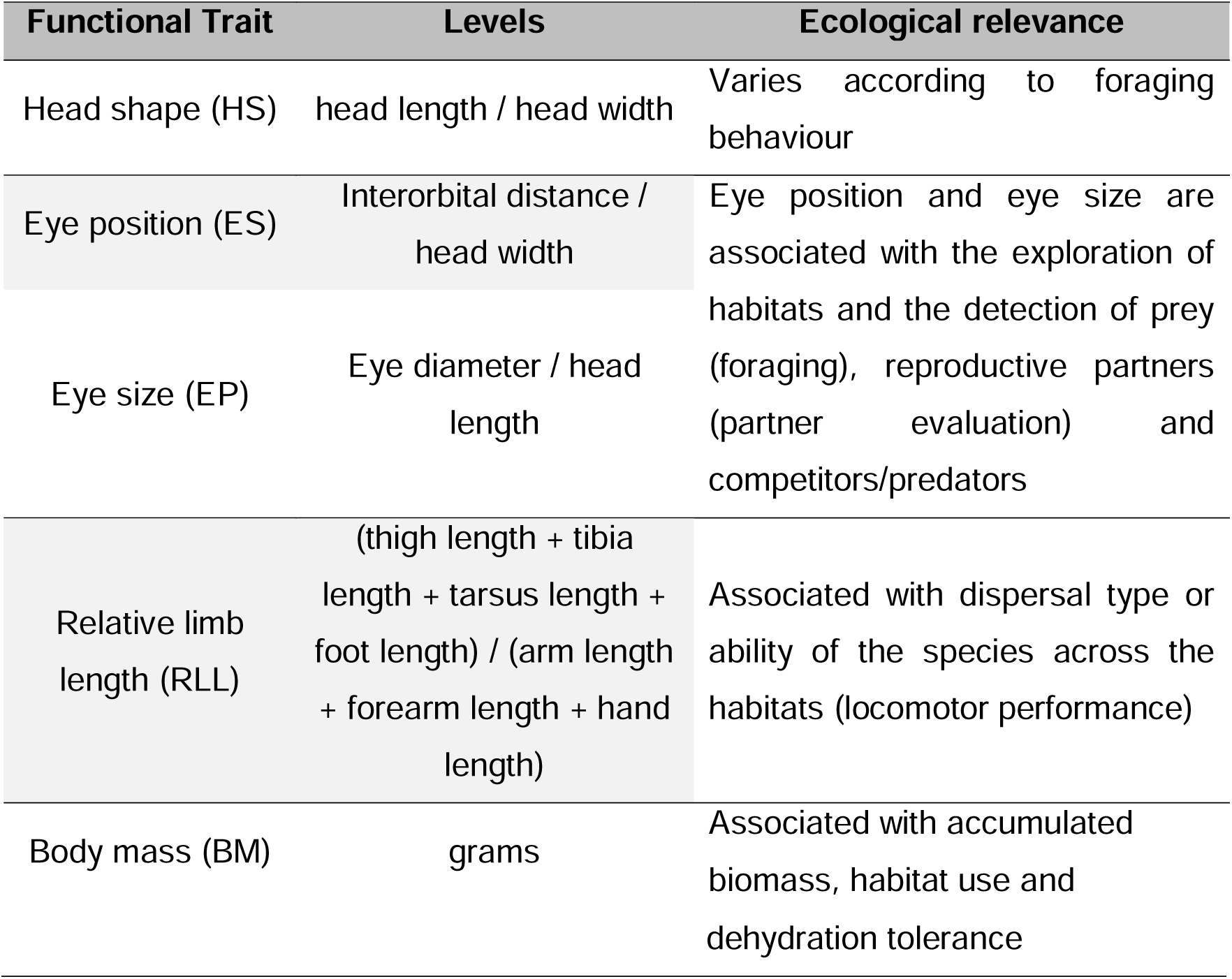
Functional traits measured (in adults).

**Figure 2:**
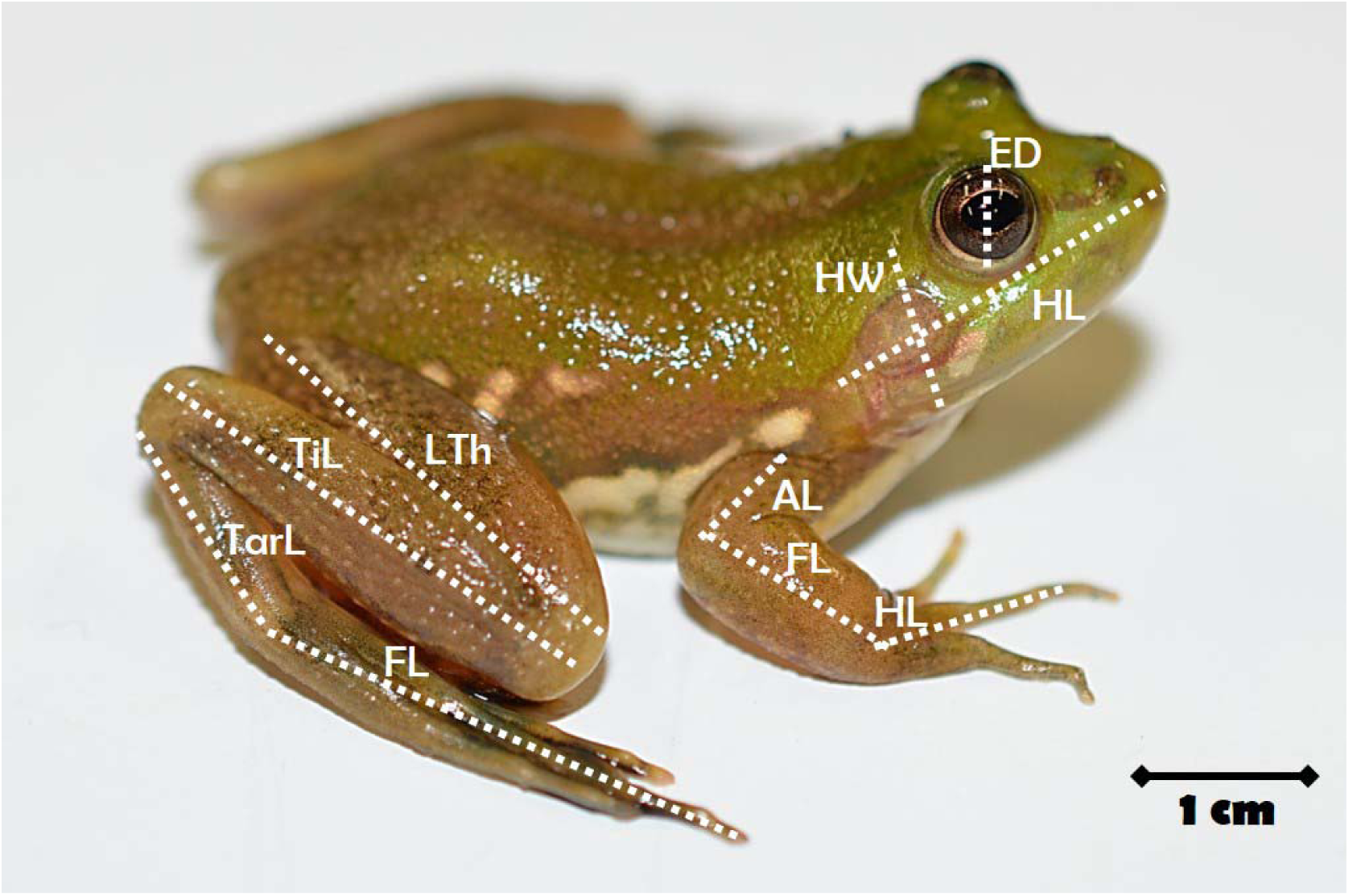
Metrics evaluated in adults (except body mass): length of thigh (LTh); tibia length (TiL); tarsus length (TarL); foot length (FL); arm length (AL); forearm length (FL); hand length (HL). In this picture: a male of *Pseudis minuta* (Amphibia, Hylidae) captured at the Parque Nacional da Lagoa do Peixe, Brazil.

### 2.4 Environmental predictors

The environmental descriptors measured in this study and their description are presented in Table S1 of supplementary material. We measured the local descriptors in the field through visual interpretation in an area around 5 meters from the edge of the ponds. We assessed area, depth and number of vegetation types around the pond and inside the pond (pond vegetation), margin configuration and type of substrate of the pond. We obtained the distance to the nearest forest fragment and the distance to the nearest sampled pond from high-resolution aerial photographs examined immediately after sampling, available from Google Earth (http://earth.google.com/), combined with field inspection.

### 2.5 Data analysis

#### 2.5.1 Partitioning anuran intraspecific trait variation across nested scales

To quantify the extent of intraspecific vs. interspecific variation of anuran functional traits along the metacommunity, we performed a variance partitioning analysis using the function “partvar” of package cati in R software version 3.4.4 (R Core Team, 2018). This function runs a variance partitioning across nested scales using the decomposition of variance on restricted maximum likelihood (REML) method (lme function), to assess variation in traits at different taxonomic levels (i.e., within-species, species and genus) and between sites (populations; Messier et al., 2010).

#### 2.5.2 Decomposing the variance in community trait composition

Following the approach proposed by Leps et al. (2011) and updated by Taudiere and Violle (2016), we decomposed the variation in community trait composition into three sources: 1) intraspecific trait variability (ITV), 2) variability due to species turnover and 3) their covariation. According to Lepš et al. (2011), trait variation will be explained by species turnover when differences in trait averages based on fixed trait values are found between habitats. On the other hand, differences in averages based on habitat specific values (e.g. ‘specific averages’) may be caused either by differences in species composition (species turnover) or intraspecific trait variation (or both). Finally, differences between specific and fixed averages may be due solely to intraspecific trait variation. These analyses were carried out using the function “decompCTRE” of the *cati* package in R software version 3.4.4 (Taudiere and Violle, 2016).

#### 2.5.3 Evaluating trait-environment relationships

We used Linear Mixed-Effect Models (LMM) to model the relationship between functional traits and local environmental predictors. This approach explicitly models the relationship within the data set using random effects or latent random variables (Breslow & Clayton, 1993; Zhang et al., 2012). We built all possible models so that all combinations between environmental predictors were evaluated. We included pond as random effect. Model selection was done using the Akaike Information Criteria corrected for small samples (AICc) to select the best model containing most information among all candidate hypotheses (Buham & Anderson, 2002). We also took into account the AICc weights (w) indicating the empirical support for each model relative to the others in the candidate set. Finally, we applied a threshold of 2 AICc units to define model support (in other words, we considered models with ΔAIC<2 as equivalent; Zuur et al., 2009). LMM were built using the ‘nlme’, ‘MuMIn’ and ‘lme4’ packages in R software version 3.4.4.

## 3 RESULTS

We registered adults of 11 species belonging to three families (Bufonidae, Hylidae and Leptodactylidae) during our sampling. The most frequent species were *Dendropsophus sanborni* and *Pseudis minuta*, occurring in 19 of the 33 sampled ponds. *Physalaemus biligonigerus, Rhinella dorbignyi* and *Scinax fuscovarius* were rare and we were unable to capture five individuals of those species. We therefore did not include them in the subsequent analyses. In Table 2 we present all species evaluated and their groupings according to their preferred habitat.

**Table 2.** Species of amphibians evaluated for intraspecific trait variation and their respective grouping based on preferred habitat.

| Groups | Species | Total Abundance in ponds |
| --- | --- | --- |
| Aquatic-terrestrial | *Leptodactylus latrans* | 52 |
|  | *Physalaemus gracilis* | 169 |
|  | *Pseudis minuta* | 186 |
|  | *Pseudopaludicola falcipes* | 153 |
| Arboreal | *Dendropsophus minutus* | 53 |
|  | *Dendropsophus sanborni* | 173 |
|  | *Hypsiboas pulchellus* | 111 |
|  | *Scinax squalirostris* | 69 |

**Table 2.**
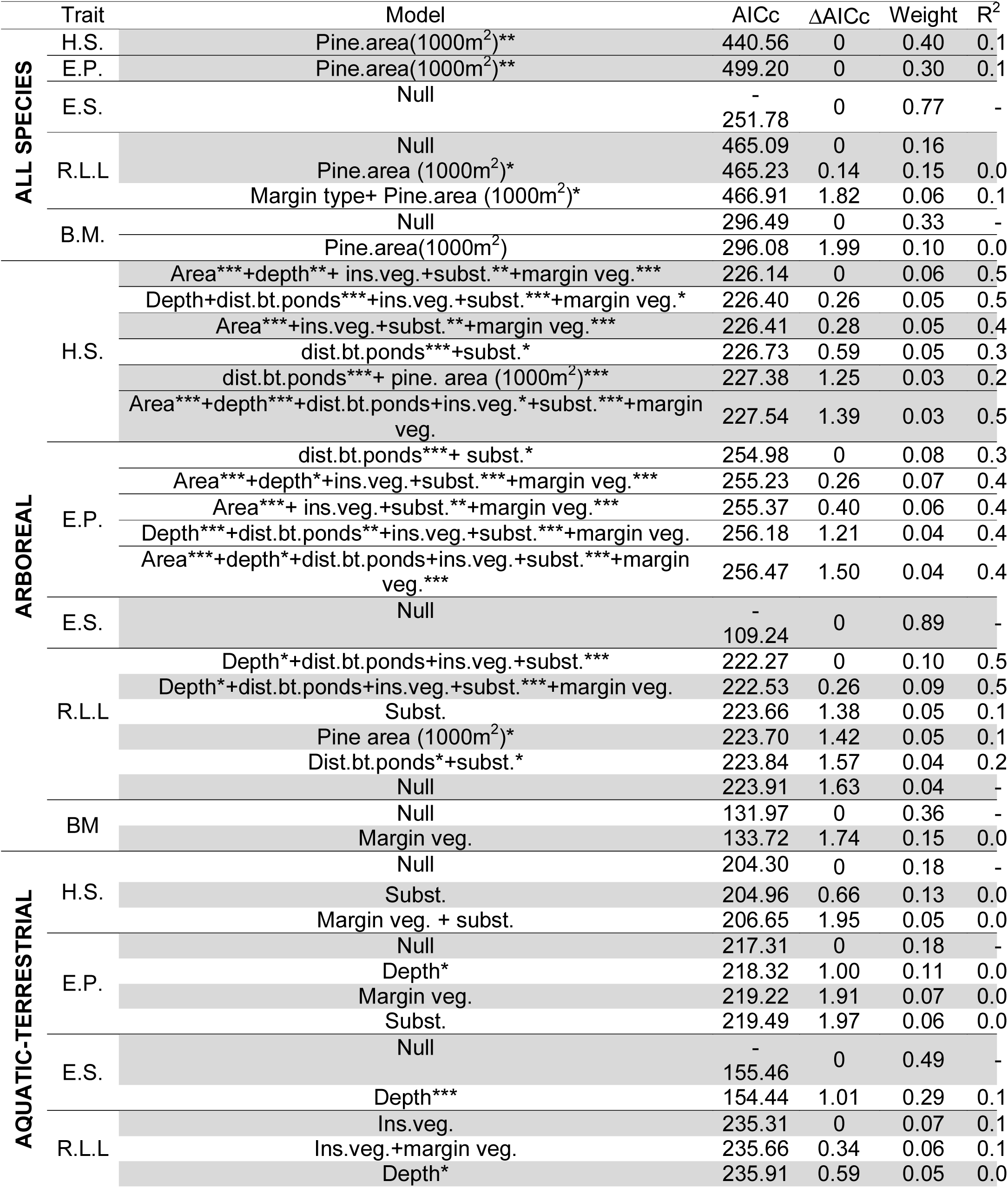
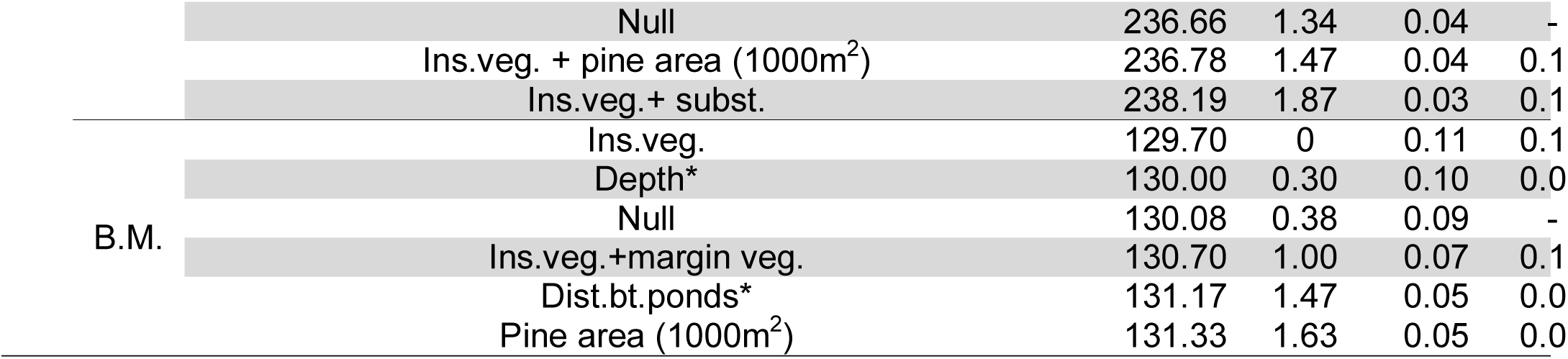
Summary of the best-adjusted models resulting from the LMM for each functional trai sampled pond metacommunity including models for all species, arboreal species set and aquatic-t species set. R^2^m (fixed effects); R^2^C (fixed + random effects). (*) *p*< .05; (**) *p*< .01; (***) *p*< .001.

### 3.1 Partitioning of anuran intraspecific trait variation across nested scales

The within-species scale accounted for most variation in head form (39%), eye size (44%) and eye position (42%) for the whole species set, while species and genus scales accounted for most variation in relative limb length (39% and 42%, respectively) and body mass (39% and 42%, respectively; Figure 3c).

**Figure 3.**
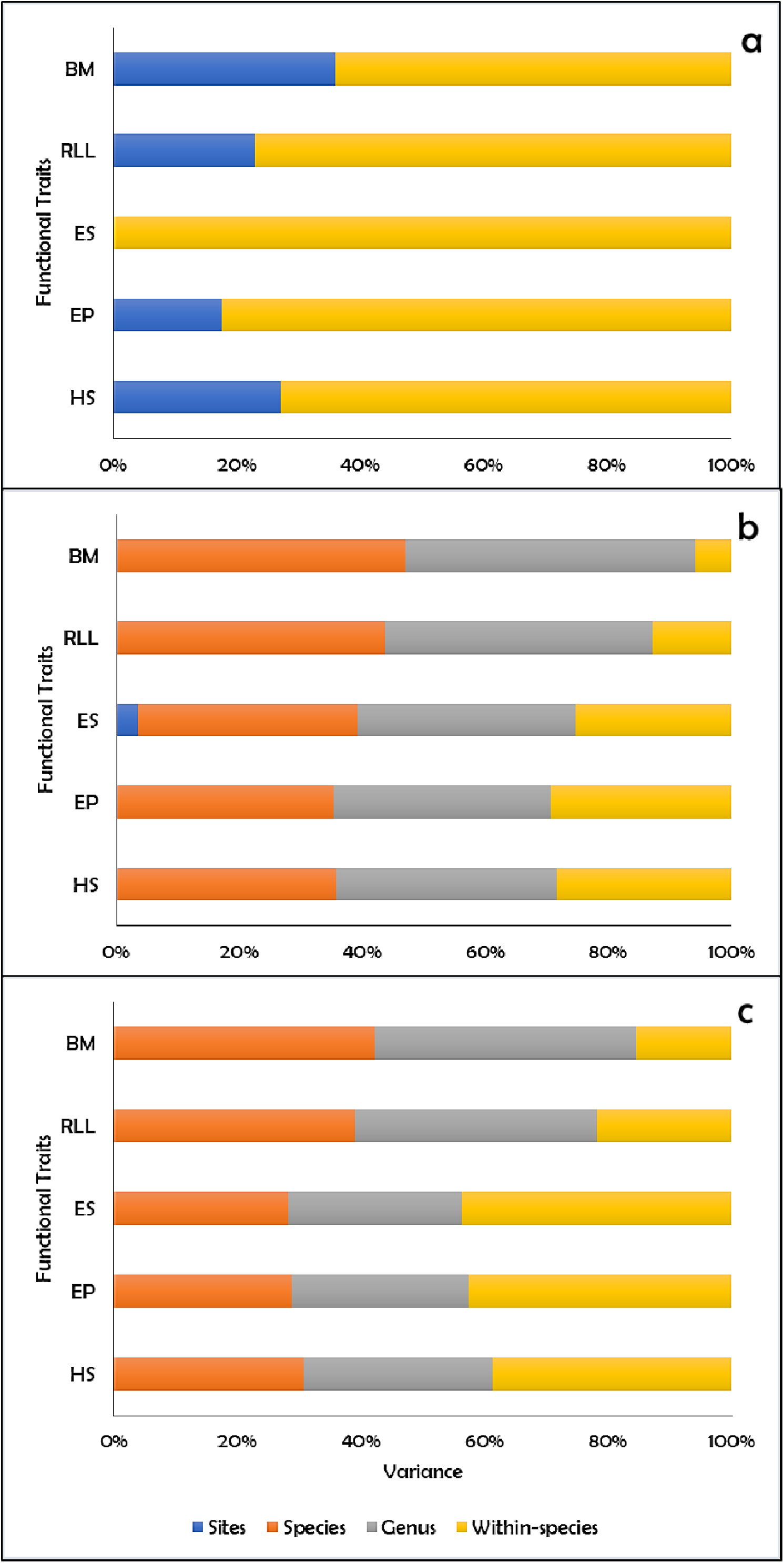
Variance partitioning of community traits for the sampled pond metacommunity: (a) arboreal, (b) aquatic-terrestrial, and (c) all species. Total between-plot variance in morphological trait values was partitioned into species turnover, intraspecific trait variation (ITV), and covariation between those components. Variance partitioning was conducted for five functional traits: head shape (HS), eye position (EP), eye size (ES), relative limb length (RLL) and body mass (BM).

We found discrepant patterns between groups. The within-species scale held 99% of variance of all traits for the arboreal species (Figure 3a), while species and genus scales accounted for the most variation of all traits for the aquatic-terrestrial species (from 75% to 95%; Figure 3b).

### 3.2 Relative contribution of species turnover and intraspecific trait variation

The relative contributions of species turnover and intraspecific trait variation (ITV) to shifts in community mean trait values in the anuran metacommunity was similar between traits, but differed between sets of species. For the entire species set the relative contribution of ITV ranged from 10% to 25%, while species turnover ranged from 56% to 88% (figure 4c). ITV and species turnover were positively correlated for all traits (figure 4c).

**Figure 4.**
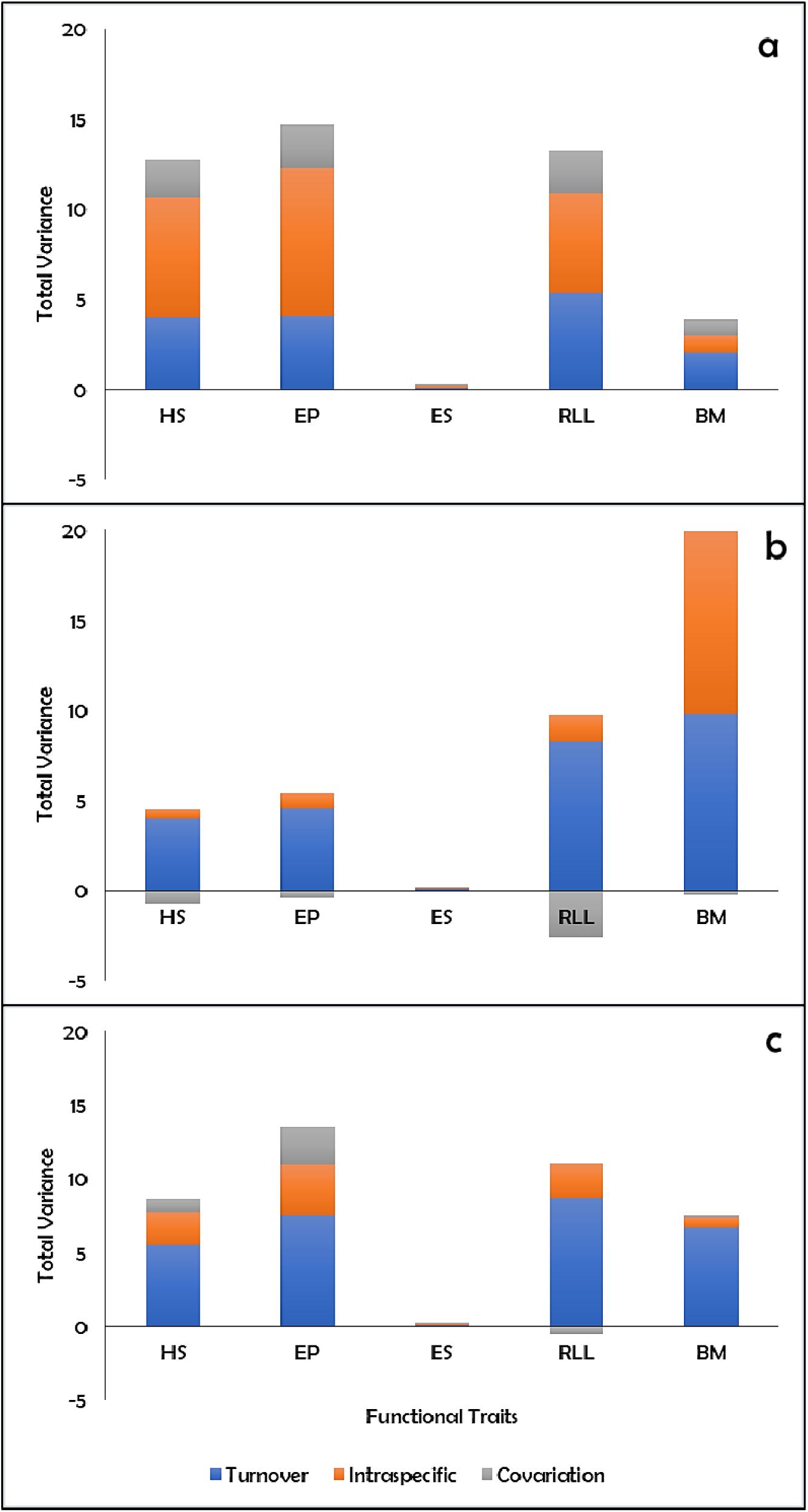
Decomposition of variance of community trait for the sampled pond metacommunity: (a) arboreal, (b) aquatic-terrestrial, and (c) all species. The total between-plot variance in functional traits was partitioned into species turnover, intraspecific trait variation (ITV), and covariation between those components. Variance partitioning was carried out for five functional traits: head shape (HS), eye position (EP), eye size (ES), relative limb length (RLL) and body mass (BM).

For the arboreal species set, the contribution of ITV was very similar to that of species turnover: the relative contribution of ITV ranged from 24% to 56%, while species turnover ranged from 27% to 52% (figure 4a). However, for the aquatic-terrestrial species set turnover was the main process governing the intraspecific trait variation and ranged from 69% to 116% (the lower value belonging to RLL, which presented a negative covariation between ITV and turnover), while ITV ranged from 11% to 31% (figure 4b).

ITV and species turnover were positively correlated for all traits in the arboreal species set (figure 4a), while this correlation was negative for the aquatic-terrestrial species set (figure 4a).

### 3.3 Trait-environment relationships

A majority of the total variance explained by fixed effects was found at groups levels (arboreal species; Table 2). In general, the null model was among the candidate models selected for the largest part of the functional traits in all sets of data evaluated and was, in some cases, the best fitted. For the whole species set, pine area presented positive and significant effects on the observed variation in head form (*F*=7.24), relative eye position (*F*=6.23) and relative limb length (*F*=3.2).

For the arboreal species set, all environmental predictors showed positive effects on the intraspecific variation of at least one of the evaluated functional traits, with the exception of margin configuration. Head form responded mostly to vegetation type (*F*=23.57), eye position to distance between ponds (*F*=12.72) and relative limb length to substrate type (*F*= 10.19). Finally, depth was the only variable influencing trait variation in the aquatic-terrestrial species set (EP: *F*= 3.49; ES: *F*=7.57; RLL: *F*=4.61; BM: *F*=4.96), and the relation was always negative, except for body mass. We did not find any relationship between head shape and the set of environmental predictors for this group. The summary tables of significance tests and all models tested are presented in supplementary material (respectively, Tables S2 and S3-S16).

**Table 3.** Summary of the most parsimonious linear mixed-effect models for each functional trait for the sampled pond metacommunity including models for all species, arboreal species set and aquatic-terrestrial species set (please see the the Table S1 for more details about the local environmental predictors). R^2^m (fixed effects); R^2^C (fixed + random effects). (*) *p*< .05; (**) *p*< .01; (***) *p*< .001.

## 4 DISCUSSION

Patterns of trait variation in anuran metacommunities and their relation with local environmental predictors are complex. ITV is generally the dominant process driving trait variation in arboreal species, species turnover seems to be the driving force behind trait variation in aquatic-terrestrial anurans. Looking into the whole set of species, we can see that turnover is, once again, the dominant process. This pattern may be explained by the fact that the whole set joins taxa with very distinct evolutionary histories, potentially responding very differently to the evaluated environmental predictors (Marques and Nomura, 2015). Also, interspecific variation represented between 60% and 90% of trait variability for the whole species set and for the aquatic-terrestrial species set, while ITV explained up to 100% of the total trait variation in arboreal anurans. Our data also points to significant effects of environmental filtering in the shaping of the trait structure in anuran metacommunities, but again with distinct factors acting depending on the preferred habitat of the species and the analysed functional trait.

### 4.1 The balance between intraspecific and interspecific trait variability

The relative importance of ITV and turnover in trait variability seems to be dependent on the preferred habitat of the species or species group under analysis, as well as the analysed trait. In general, we found a greater variation in the scale between species and a substantial contribution of turnover for most of the traits of aquatic-terrestrial species group and for the total set of species. For the neotropical anuran metacommunity in our study, as well as for several groups of organisms, species turnover seems to be the dominant process behind trait variation (e.g. Siefert et al., 2015 Derroire et al., 2018). Species turnover tends to be stronger at larger spatial scales, where environmental heterogeneity is patent, whereas phenotypic adjustments should be dominant at small spatial scales and within short time frames (Jung et al., 2010; Kichenin et al., 2013; Kumordzi et al., 2015; Chalmandrier et al., 2017).

From an evolutionary perspective, ITV within and between populations is influenced by physiological, ontogenetic, genetic and ecological factors (Miller and Rudolf 2011; Ingram and Shurin 2009; Zhao et al., 2014). Genetic and ecological factors have been pointed out as the main drivers of trait variation in anuran populations and between ecomorphological guilds (e.g. Lampert et al., 2003; Trakimas et al., 2003; Zeisset and Beebee, 2003; Palo et al., 2004; Telles, 2005); this is due – at least partially – to the fact that they are easier to evaluate than physiological and ontogenetic factors *per se*, considering that these should be closely correlated, influencing and being influenced by both genetic and ecological contexts.

Anurans present high phenotypic plasticity in response to environmental conditions (Michel 2011; Marques and Nomura, 2015), which may be higher for some traits, as detected in our study. Terrestrial and arboreal species are subject to distinct microenvironmental conditions, and they may differ considerably in their foraging strategies and habitat use. It should not thus come as a surprise that arboreal and the aquatic-terrestrial anurans show distinct ITV patterns.

Head shape, eye size and eye position presented significant variation within species, contrarily to our predictions. Usually, head and eye characteristics contribute to significant functional dissimilarity between anurans, but most of the arboreal species sampled here seem to be quite redundant in relation to these traits. So, most of the variation turned out to be intraspecific, in agreement with findings for other anuran populations which also showed high functional overlap in these traits (Silva et al., 2008). ITV dominance has been observed for top predators, including between individuals of the same ontogenetic classes or age cohorts (Zhao et al., 2014). If local conditions favour genetic diversification (Silva et al., 2008; Vieira et al., 2008), one would expect within-population variation in functional traits. In fact, significant within-population allelic diversity has already been reported for anuran populations of the Cerrado and Atlantic Forest biomes (Silva et al., 2008; Vieira et al., 2008), potentially contributing to the co-occurrence of phenotypically distinct individuals within populations.

On the other hand, body mass and relative limb length tended to vary more between species and genera than within populations, suggesting that analyses of the functional structure of these attributes may be measured through mean values per species (Garnier et al., 2001; Griffiths et al., 2016). Body mass and relative limb length are strongly related to the rates of dispersion and habitat use; discrepancies in allometric relationships between arboreal and terrestrial species relating to body mass seem to be related to the differential effect that gravity exerts on species in aquatic and terrestrial environments, promoting adequate body mass adjustments for each of these environments (Vidal-García and Keogh, 2015). However, ITV assessments for body mass and relative limb length should not be disregarded, as both high intra and interspecific variation across geographical and temporal scales have been reported for these traits (e.g. Dodd, 2010; Santini et al., 2018; Amado et al., 2019). For example, body mass reflects physiological states and water and heat retention abilities (Wells, 2010; Dodd, 2010) and is often used as proxy for the risk of extinction (Purvis et al., 2000; Cardillo et al., 2005). Because extinction acts on populations, intra and inter-population variation in traits that mirror individual fitness and population health may be important indicators of the extinction risk of the populations under study.

### 4.2 Complex relations between traits and environment in anurans

As predicted, trait-environment relationships were more evident within species sets that share similar habitat preferences than for the total set of species. Overall, for several traits, ITV responded to more than one environmental predictor, showing the complex nature of trait-environment relationships. Pond configuration – particularly area, depth, types of substrate and margin vegetation, as well as the distance between ponds – was the main driver of ITV. All these reflect environmental heterogeneity and tend to vary greatly in narrow space scales and short time frames, probably contributing to the high phenotypic plasticity observed (Welborn et al., 1996; Werner et al., 2007; Both et al., 2011). Depth, for instance, was significantly associated with all the evaluated functional traits and, together with pond area, is a proxy for hydroperiod and evapotranspiration rates of ponds (Welborn et al., 1996; Williams, 2006; Werner et al., 2007). Changes in pond depth and area directly affect the reproductive biology and the physiological processes associated with water conservation in anurans (Wells, 2010; Amado et al., 2019). Again, this is connected to the observed variation in body mass between species and genera. Here, deeper ponds harboured, in average, weightier aquatic and terrestrial species. This reflects the fact that larger bodies (by mass) allow for higher water and heat retention, which should be particularly critical in species that are in constant contact with the substrate, as is the case of aquatic and terrestrial anurans (Wells, 2010; Amado et al., 2019).

Other environmental aspects may also influence the ability of anurans to regulate body temperature and osmotic rate, such as the percentage of humidity in the soil around the ponds, as well as pond vegetation (Wells 2007; Olalla-Tárraga and Rodríguez, 2007; Gouveia & Correia, 2016). Behavioural responses, like the use of particular climatic microhabitats – such as those created in humid soils or within semi-aquatic vegetation –, are well known in anurans (Wells 2007, Vitt and Caldwell 2009) and tend to be far more frequent in species and individuals with smaller body sizes or occurring in colder regions (Amado et al., 2019). In addition, variation in individual size is expected to be closely related to resource availability (“resource rule” - McNab, 2010), which is less variable in larger and deeper environments.

In theory, all dimensions of anuran diversity in ponds gradually increase with structural complexity, but decay with distance between ponds and between ponds and forest fragments (e.g. Hill et al., 2017). In fact, increased diversity is driven by the number of niches available, which tends to be higher in complex environments (e.g. Ruhi et al., 2014; Heino et al., 2015; Spasojevic et al., 2016), while limited dispersal ability imposed by limb morphology and physiological restrictions (risk of desiccation) compromise long-distance anuran dispersal (Wells, 2010; Amado et al., 2019). Differences in life history, morphology, and locomotion mode directly affect trait-environment relationships and species turnover dominance (Fabrezi et al., 2017), as observed in our study. According to our predictions, distance between ponds mostly influenced variation patterns in traits associated with habitat exploitation in arboreal species – head shape, eye position and, particularly, limb length. Arboreal species use forest fragments that may often be distant from the ponds and move to reproductive sites through large leaps (da Silva et al., 2012). In contrast, aquatic and terrestrial species have relatively smaller limbs compared to arboreal species, so are more limited in terms of long-distance dispersal, but on the other hand are better adapted to swimming. They use burrows or any type of substrate to take refuge during the day (e.g. leptodactylides).

Trait variation for the whole species set was strongly associated to the area of *Pinus* around ponds (1000 m^2^ buffer). Interestingly, the relative area of *Pinus* plantations within that buffer seems to contribute positively to the variation of the traits related to habitat exploitation. *Pinus* trees acidify wetlands, change soil conditions, and create new shaded areas in ponds and adjacent areas (Whatling et al., 2011; Machado et al., 2012; Saccol et al., 2017). In combination, these environmental modifications probably impose physiological and behavioural barriers (e.g. thermoregulation and displacement, Rothermel and Semlitsch, 2002; Parris, 2004; Saccol et al., 2017). As a consequence, population resilience in these environments may be conditioned by the presence of high rates of variation of these attributes (phenotypic plasticity, Werner et al., 2007, Michel, 2011). Nonetheless, this result should be taken with precaution, because while trait variation may be increasing, taxonomic diversity may decrease in the long run. Indeed, herpetological diversity is known to decrease in areas altered by silviculture in southern Brazil (e.g. Machado et al., 2012; Saccol et al., 2017) and in areas across the globe (e.g. Popescu and Hunter Jr., 2011). The presence of exotic plantations affects the local conditions of the ponds, leading to biotic homogenization between environments and consequently reducing functional and taxonomic diversity on the long-term (Machado et al., 2012; Saccol et al., 2017).

## 5 CONCLUSIONS

Functional traits related to arboreal exploitation in adult anurans (HS, ES and EP) were more variable on the intraspecific level, which may be related to individual adaptations to local conditions (e.g. microhabitat). In contrast, interspecific variation was dominant in traits related to horizontal movement and foraging of adult anurans (BM and RLL). This may be due to differences in dispersal mode and ability, as well as to differences in the species’ trophic levels. These may reflect solutions to reduce overlap in the trophic space, facilitating coexistence between species (Cloyed and Eason, 2017). We stress the importance of considering patterns of intraspecific variation in functional traits for the evaluation of the functional structure of metacommunities. Also, ecological differences between species may also influence the outcomes of analyses of ITV versus turnover; thus, analyses should focus on the whole set of species under analysis but also distinct functional guilds.

## Author contributions

All authors contributed to the study conception and design. Material preparation, data collection and analysis were performed by Diego Anderson Dalmolin and Maria João Ramos Pereira. The first draft of the manuscript was written by Diego Anderson Dalmolin and all authors commented on previous versions of the manuscript. All authors read and approved the final manuscript.

